# Neonatal T cell maternal microchimerism is common and associates with early pregnancy SARS2 vaccination

**DOI:** 10.64898/2026.10.23.761412

**Authors:** Jacquelyn Weaver, John Houck, Yonghou Jiang, Cara Tobey, Jia Wong, Swati Shree, Whitney E Harrington

## Abstract

The fetus acquires a graft of maternal cells in utero, known as maternal microchimerism (MMc), which may be enriched for T cells. However, little is known regarding the determinants of T cell trafficking into the fetus or regarding the prevalence of T cell MMc in healthy neonates. Using a closely followed cohort, we determined the association between maternal vaccination and acute infection and the detection of T cell MMc in cord blood, as measured using polymorphism-specific qPCR. We identified an association between early pregnancy SARS2 mRNA vaccination, a potent T cell stimulus, and increased detection of T cell MMc.

## INTRODUCTION

During pregnancy, rare maternal cells traffic to the fetus, a phenomenon known as maternal microchimerism (**MMc**) (1). In cord blood, we have identified MMc in many immune lineages and the highest level in the T cell compartment (2). Detection of MMc in cord blood associates with protection from symptomatic malaria, non-malarial fever, and respiratory tract infection in the offspring (3, 4). Further, we have identified that birth MMc associates with accentuated responses to BCG vaccination in South African infants (5). These data are supported by murine models demonstrating that T cell MMc may both directly and indirectly influence offspring protection from infection (6, 7). Together these observations suggest that MMc, particularly T cell MMc, may play an underrecognized role in shaping fetal and infant immunity.

Relatively little is known about the factors that determine the level and distribution of MMc in the fetus and neonate, although we have identified that maternal malaria in the first half of pregnancy (8) and recently diagnosed HIV (5) are associated with level of MMc in the neonate. These observations, as well as earlier work demonstrating MMc in the fetus at the beginning of the second trimester (9), suggest that maternal immune activation early in pregnancy may be a key determinant of MMc trafficking. Indeed, activated T cells are known to upregulate tissue trafficking molecules (10), which may facilitate their movement across the placenta into the fetal circulation.

Based on these data, we hypothesized that in a United States cohort maternal immune stimuli in the first half of pregnancy, including vaccination and infection, would associate with increased T cell MMc in the neonate.

## METHODS

### Cohort

Individuals aged 18-45 with singleton pregnancies receiving obstetric care at the University of Washington affiliated clinics were approached for enrollment into the Fetal Immune Development Study at the time of their first prenatal visit between 2022-2023. Women with multifetal gestation, fetal aneuploidy, or immunodeficiency or modulation were ineligible. Peripheral blood mononuclear cells (**PBMC**) were isolated from maternal whole blood collected into ACD tubes during routine prenatal visits (8-12 gestational weeks (**gw**), 24-28 gw) and at the time of delivery. Cord blood was collected into ACD tubes via cross-clamping, cleaning, and direct cannulation of the cord, and cord blood mononuclear cells (**CBMC**) were isolated. Women received a Redcap survey at the end of each trimester querying any acute illnesses (influenza, SARS2, other respiratory, or gastrointestinal) or any vaccines (influenza, SARS2, Tdap, or RSV) received over the prior 14 weeks. Clinical data including maternal age, pregnancy history, prior and current health conditions, vaccinations received, and medications were abstracted from the medical chart following delivery. Reported timing of vaccinations (all) and illnesses (where reported) were verified with the medical record. All participants provided written informed consent, and the study was approved by the UW IRB (STUDY00001636, STUDY00004878).

### Enrichment of T cells from cord blood mononuclear cells

T cells were enriched from CBMC using magnetic-activated cell sorting (**MACS**). Briefly, ∼10 million CBMC were thawed and stained with PE-conjugated anti-human CD3 antibody (BioLegend) for 20 min at 4°C in the dark. Following staining, cells were washed with MACS buffer (PBS with 2% FBS and 2 mM EDTA), resuspended, and 20µL of Anti-PE MicroBeads UltraPure (Miltenyi Biotec) was added. The cell suspension was mixed and incubated for 20 min at 4°C. The cells were washed and resuspended in 500µL of buffer for magnetic separation. MS columns were placed in the magnetic field of a MACS Separator and rinsed with 500 µL of buffer before the cell suspension was applied. The unbound cells in the flowthrough were designated as non-T cells. The columns were then removed from the separator and placed on collection tubes. The magnetically enriched T cells were then eluted in 1mL of MACS buffer.

### Measurement of maternal microchimerism

Genomic DNA was extracted from the enriched T cells using the PrepFiler™ Forensic DNA Extraction Kit (Thermofisher) with modifications to improve recovery including the addition of carrier RNA (Qiagen) and an additional elution. MMc was detected and quantified using a set of recently developed qPCR assays targeting deletion/insertion polymorphisms (*Houck, BioRxiv, 2026*). Briefly, mothers and infants were genotyped at 18 loci to identify a non-inherited allele specific to each mother in the background of her infant. The informative allele was then targeted using a highly sensitive and specific qPCR assay. The assessment of MMc in each infant included standard curves for the maternal target as well as betaglobin, such that a numerator (MMc genomic equivalents (**GE**)) and a denominator (infant GE) could be calculated, and MMc could be expressed as detected / not detected or as a quantitative value.

### Statistical analysis

Immune stimuli across pregnancy were modeled to predict detectable T cell MMc (0/1) using logistic regression adjusting for the GE tested for each infant. We first considered any exposure to the stimuli (e.g., any SARS2 mRNA vaccination), and then in a second model included exposure in early pregnancy (2-19.9 weeks gestation) or late pregnancy (20-42 weeks gestation) as separate predictors (e.g., SARS2 mRNA vaccination in early pregnancy and/or late pregnancy). RSV vaccination was modeled as a single predictor as all exposure occurred in late pregnancy. The impact of Tdap vaccination was not modeled as 95% of participants received this during pregnancy. The impact of influenza infection could not be modeled with logistic regression because all neonates of women with infection had detectable T cell MMc (the outcome did not vary). Predictors were modeled together in fully adjusted models, and all models used robust standard errors. The distribution of MMc levels (0, low: 1-100 per 10^6^, high: >100 per 10^6^) was assessed using χ^2^ tests.

## RESULTS

### T cell microchimerism is common

We followed 63 women from enrollment through delivery with complete collection of exposure data as well as delivery samples. Women were a median age of 34 (range: 23-44) and were predominantly multigravid (**Table S1**). Median gestational age at delivery was 39 gw (range: 35-42 gw). Infants were evenly split female (49%) and male (51%) and were delivered via vaginal birth (62%), labored Cesarean (C)-section (25%), or scheduled C-section (13%). A median of 83,787 (range: 4,385-232,893) infant GE were tested for T cell MMc. The majority of infants (70%) had detectable T cell MMc, with a median level of 45 (range: 5-30,884) per 10^6^ infant GE. None of the above noted characteristics in the mother or the infant were associated with detection of T cell MMc.

### Immune stimuli are frequent and distributed across pregnancy

Maternal vaccination was common with 60 (95%) women reporting Tdap vaccination, 34 (54%) influenza vaccination, 23 (37%) SARS2 vaccination, and 15 (24%) RSV vaccination (**Table 1**). Influenza and SARS2 vaccination were evenly split between early and late pregnancy (**Table 1, Figure S1**). As expected, Tdap and RSV vaccination were exclusively received during late pregnancy. Because pregnancy could span multiple vaccine seasons, one woman reported two influenza vaccines and a second two SARS2 vaccines. All women reporting SARS2 vaccination received an mRNA-based vaccine. RSV vaccination receipt was relatively low as it was only available during the final year of the cohort.

**Table 1.**
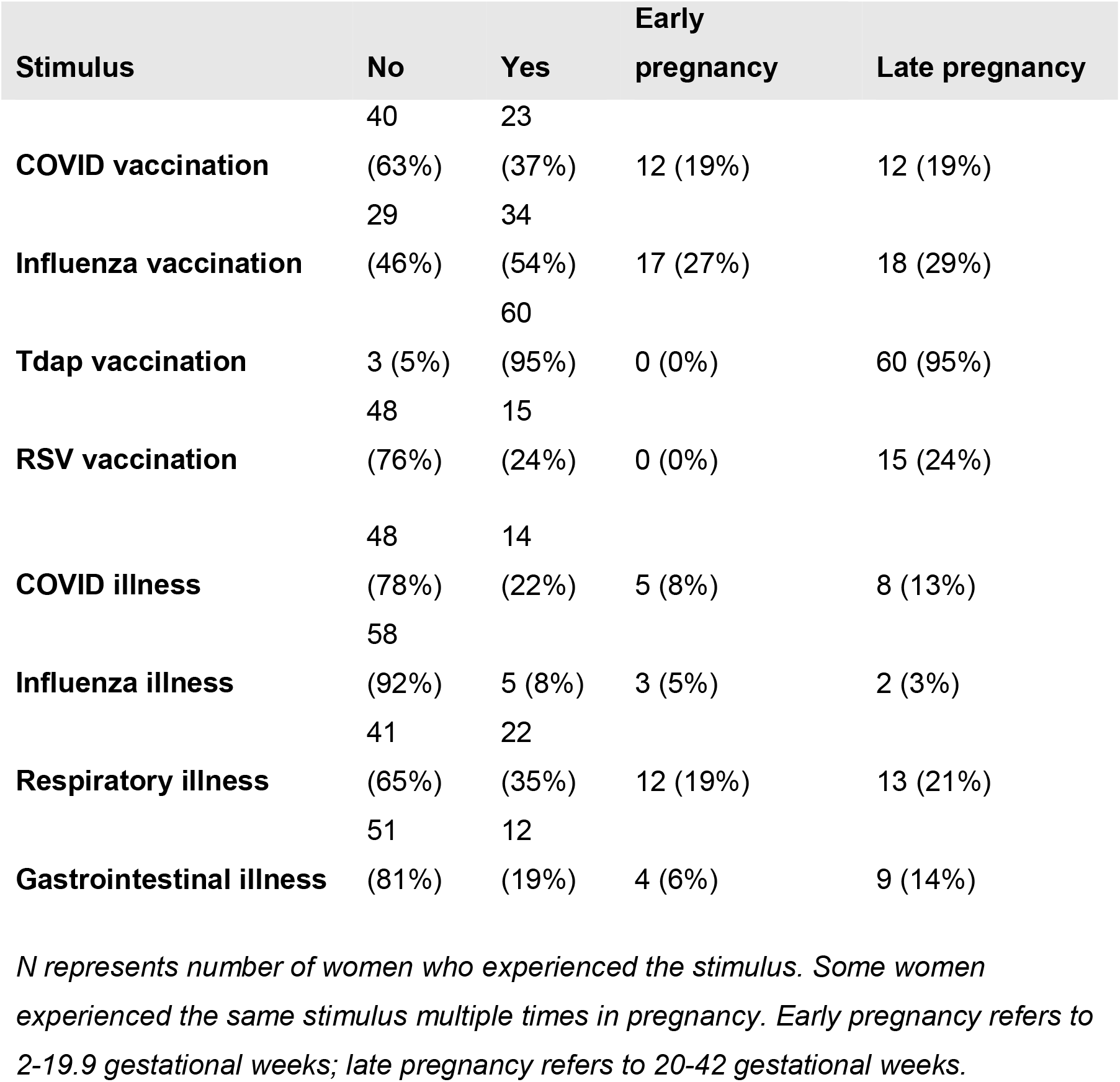
Immune stimuli experienced during pregnancy.

Fewer women reported acute infection, with 22 (35%) reporting respiratory illness, 14 (22%) SARS2 infection, 12 (19%) gastrointestinal illness, and 5 (8%) influenza infection. Like vaccination, the distribution of infection was similar in early and late pregnancy (**Table 1**). One woman reported three respiratory tract infections, six women reported two respiratory tract infections, and two women reported two gastrointestinal infections.

### SARS2 vaccination in early pregnancy associates with T cell MMc

When any vaccination across pregnancy was considered, SARS2 mRNA vaccination (AOR=4.45, p=0.08), but not influenza or RSV vaccination, was associated with a trend toward increased odds of detection of T cell MMc in the neonate (**Figure 1**). When considered by timing of vaccination, this was driven by early (AOR=9.60, p=0.05) but not late (AOR=1.78, p=0.5) pregnancy exposure. Regarding level of T cell MMc, there appeared to be a shift from no T cell MMc to low MMc associated with early SARS2 vaccination versus no vaccination (no MMc: 8% vs 35%; low: 75% vs 38%, high: 18% vs 28%, p=0.06). Infection was not associated with statistically significant association with T cell MMc; however, there were very few serious infections. When SARS2 and influenza infections were modeled as a combined outcome, there was a non-significant but positive association with early (AOR=4.38, p=0.2) but not late (AOR=1.23, p=0.8) exposure and detection of T cell MMc.

**Figure 1.**
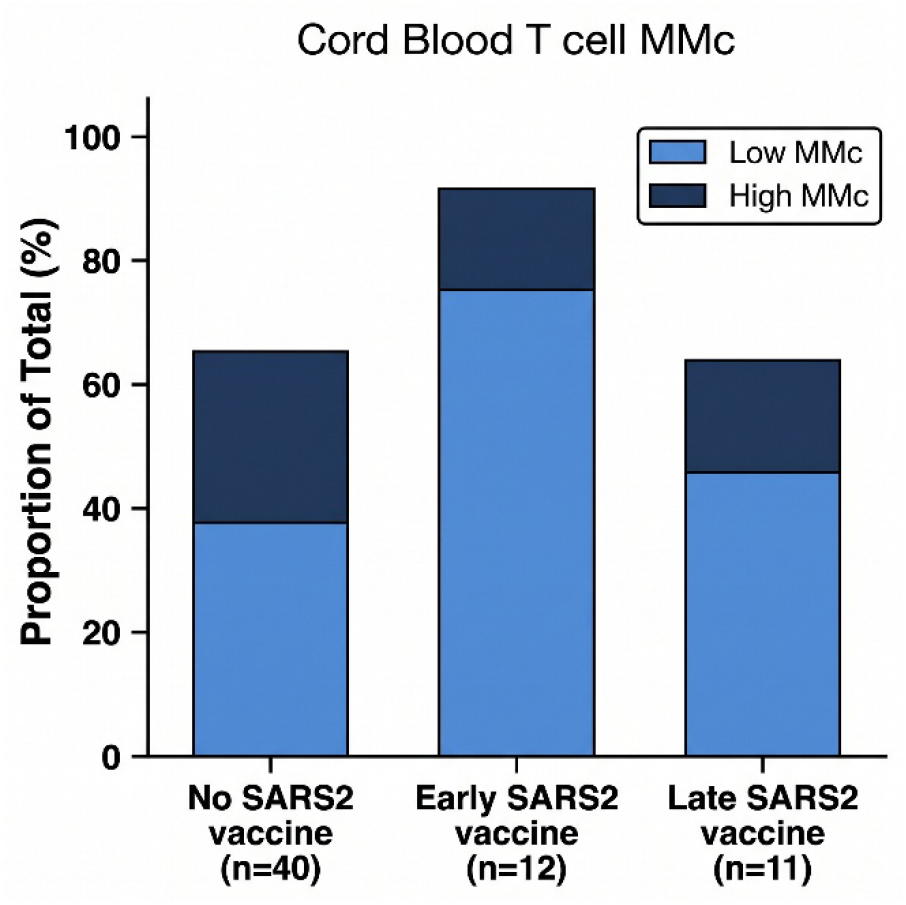
Proportion of cord blood samples positive for T cell MMc by maternal SARS2 vaccine status. T cell MMc level indicated as low (1-100 per 10^6^ infant GE) or high (>100 per 10^6^ infant GE). In adjusted logistic regression maternal SARS2 vaccination in early (AOR=9.60, p=0.05) but not late (AOR=1.78, p=0.5) pregnancy was associated with increased odds of detection of T cell MMc in cord blood. Early pregnancy: 2-19.9 gestational weeks; late pregnancy: 20-42 gestational weeks.

## DISCUSSION

The field of MMc continues to grow with accumulating evidence that these rare maternal cells acquired in utero play an outsized role in fetal and infant immune development. Here we provide evidence that the majority of healthy neonates have detectable T cell MMc in cord blood, present at a wide range of levels. Further, we identified in this United States cohort that SARS2 mRNA vaccination early in pregnancy associated with increased odds of detection of T cell MMc.

To-date, a limited set of data has characterized the distribution of MMc in immune cell subsets. Our prior work from 27 United States cord bloods identified the highest frequency of MMc in T cells; however, characteristics of the mother or the neonate were not available for analysis (2). Our findings from the present cohort are consistent with this observation, demonstrating that 70% of neonates had detectable MMc at birth. They are also consistent with our observations from South African infants during the first six months of life, in which we identified both CD4 and CD8 T cell MMc (11). Importantly, in the present study, we utilized a recently developed polymorphism-based qPCR method for the detection of MMc, which overcomes the limitations of previously utilized HLA-based approaches (5, 12), in which pairs with the highest MMc may be excluded. The converging evidence provided in these studies suggests that most offspring acquire T cells from their mothers in utero.

In the present United States-based cohort, maternal immune stimuli were limited to vaccination and self-reported infection. We identified a positive association between early pregnancy SARS2 mRNA vaccination and detection of T cell MMc. This is consistent with our prior observations of an association between early pregnancy malaria infection (8) and recently diagnosed HIV (5) and MMc in the neonate, suggesting an intimate connection between the maternal immune system in the first half of pregnancy and the trafficking of cells to the fetus. This contrasts with maternal antibodies, in which the majority are transmitted to the fetus in late pregnancy (13). SARS2 mRNA vaccination is known to be highly T cell immunogenic (14), and it may be that the degree of T cell stimulation associated with this vaccine was greater than with other maternal vaccine platforms. Lending support to this model, the combined outcome of SARS2 infection and influenza infection, both of which would be expected to potently stimulate maternal T cells, was non-significantly associated with increased detection of T cell MMc. Should these observations be confirmed, it would suggest that the transmission of MMc could be intentionally modulated through T cell-directed maternal vaccination.

Our study has a few limitations. First, the number of T cells recovered from each infant CBMC varied. Reassuringly, there was no association between increasing GE assessed and increased detection of T cell MMc in the neonate. Second, we relied on self-report of maternal illness, as most mothers do not present to care for non-serious illnesses, and we may not have captured all illnesses. Third, our cohort was relatively small. Because of high vaccination exposure in this cohort, we were able to detect the effect of maternal SARS2 vaccination. Intriguingly, there was a positive association between SARS2 and influenza infection and detection of T cell MMc; however, we lacked the power to detect this association with statistical significance.

In conclusion, utilizing a closely followed cohort, we demonstrate both a high prevalence of T cell MMc in healthy neonates and an association with early pregnancy SARS2 vaccination. Given accumulating evidence that MMc influences susceptibility to infection, future efforts should focus on the ability to modulate MMc through early pregnancy T cell-directed maternal vaccination strategies.

## ACKNOWLEDGEMENTS

We would like to acknowledge the assistance of Quinn Peters, Abigail Spray, Aiesha Abdulla, and Nolawit Mulugeta in processing samples from the FID cohort.

## USE OF ARTIFICIAL INTELLIGENCE

Artificial intelligence was used in the generation of Figure 1.

## SUPPLEMENT

**Supplemental Table 1.** Participant Characteristics.

| **Characteristic** | **N=63** |
| --- | --- |
| **Age (years)** | 34 (23,44) |
| **Parity** |  |
| Primiparous | 32 (51%) |
| Multiparous | 31 (49%) |
| **Gestational age (weeks)** | 39 (35,42) |
| **Mode of delivery** |  |
| Vaginal | 39 (62%) |
| Labored C-section | 16 (25%) |
| Scheduled C-section | 8 (13%) |
| **Fetal sex** |  |
| Female | 31 (49%) |
| Male | 32 (51%) |

*Data presented as median (range) or n (%)*

**Supplemental Figure 1.**
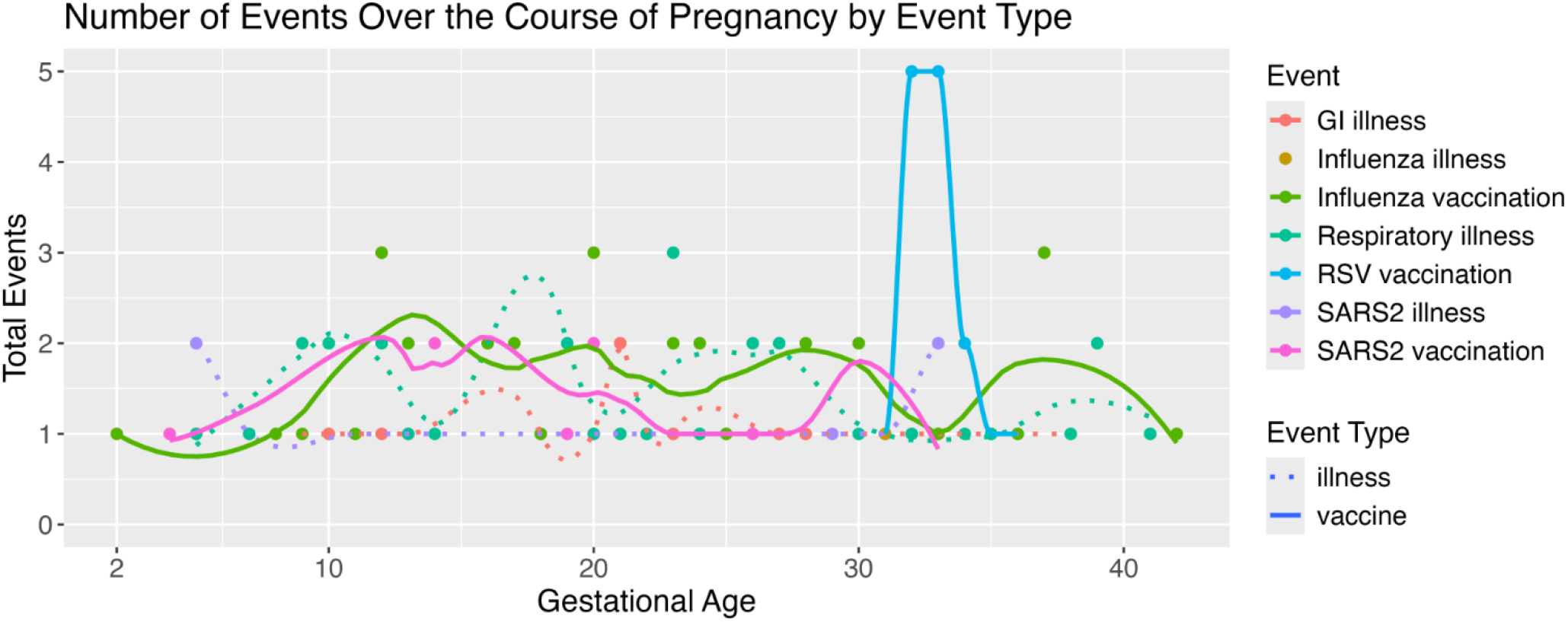
Immune stimuli including vaccination and acute illness experienced by women (n=63) across pregnancy. Data represented as individual events (dots) and LOWESS curves with span 0.35.

